# Quantitative live-cell imaging of GPCR downstream signaling dynamics

**DOI:** 10.1101/2021.08.22.457285

**Authors:** Ryosuke Tany, Yuhei Goto, Yohei Kondo, Kazuhiro Aoki

**Affiliations:** Quantitative Biology Research Group, Exploratory Research Center on Life and Living Systems (ExCELLS), National Institutes of Natural Sciences, 5-1 Higashiyama, Myodaiji-cho, Okazaki, Aichi 444-8787, Japan; Division of Quantitative Biology, National Institute for Basic Biology, National Institutes of Natural Sciences, 5-1 Higashiyama, Myodaiji-cho, Okazaki, Aichi 444-8787, Japan; Department of Basic Biology, School of Life Science, SOKENDAI (The Graduate University for Advanced Studies), 5-1 Higashiyama, Myodaiji-cho, Okazaki, Aichi 444-8787, Japan

**Keywords:** GPCR, signaling dynamics, fluorescence imaging, dopamine, serotonin

## Abstract

G-protein-coupled receptors (GPCRs) play an important role in sensing various extracellular stimuli, such as neurotransmitters, hormones, and tastants, and transducing the input information into the cell. While the human genome encodes more than 800 GPCR genes, only four Gα-proteins (Gα_s_, Gα_i/o_, Gα_q/11_, and Gα_12/13_) are known to couple with GPCRs. It remains unclear how such divergent GPCR information is translated into the downstream G-protein signaling dynamics. To answer this question, we report a live-cell fluorescence imaging system for monitoring GPCR downstream signaling dynamics at the single-cell level. Genetically encoded biosensors for cAMP, Ca^2+^, RhoA, and ERK were selected as markers for GPCR downstream signaling, and were stably expressed in HeLa cells. GPCR was further transiently overexpressed in the cells. As a proof-of-concept, we visualized GPCR signaling dynamics of 5 dopamine receptors and 12 serotonin receptors, and found heterogeneity between GPCRs and between cells. Even when the same Gα proteins were known to be coupled, the patterns of dynamics in GPCR downstream signaling, including the signal strength and duration, were substantially distinct among GPCRs. These results suggest the importance of dynamical encoding in GPCR signaling.

## INTRODUCTION

G-protein-coupled receptors (GPCRs), the largest family of membrane proteins, are triggered by various types of ligands, such as neurotransmitters (e.g., dopamine, serotonin), peptide hormones (e.g., angiotensin II), tastants, and light, to name a few. More than 800 GPCR genes are encoded in the human genome, and play indispensable roles in a vast variety of biological processes [1]. Moreover, GPCRs are one of the most important targets for drug discovery, as demonstrated by the fact that GPCRs constitute targets of approximately one-third of all Food and Drug Administration (FDA)-approved drugs [2].

Upon ligand stimulation, GPCRs activate heterotrimeric G proteins composed of alpha, beta, and gamma subunits (Gα, Gβ, and Gγ) [3]. The activated GPCRs promote guanine nucleotide exchange of Gα proteins from a GDP-bound inactive state to a GTP-bound active state [4]. The different types of GTP-loaded Gα proteins bind to and activate/inactivate different downstream effectors (**Figure 1A**). Gα_s_ positively regulates and Gα_i/o_ negatively regulates adenylate cyclases, which are enzymes that catalyze cyclic AMP (cAMP) production [5,6]. Gα_q/11_ induces intracellular Ca^2+^ increase through phospholipase C beta (PLCβ), while Gα_12/13_ activates the Rho family of small GTPases [7–9]. It has also been reported that Gα_s_, Gα_i/o_, and Gα_q/11_ regulate the ERK MAP kinase pathway [10–13]. In addition to Gα proteins, accumulating evidence supports the independent roles of the Gβγ complex in activating Rho family GTPases [14]. Other well-known mediators of GPCR signaling, including regulators of G protein signaling (RGSs), G-protein-coupled receptor kinases (GRKs), and β-arrestin, have also been shown to modulate the ERK MAP kinase pathway [10].

**Figure 1.**
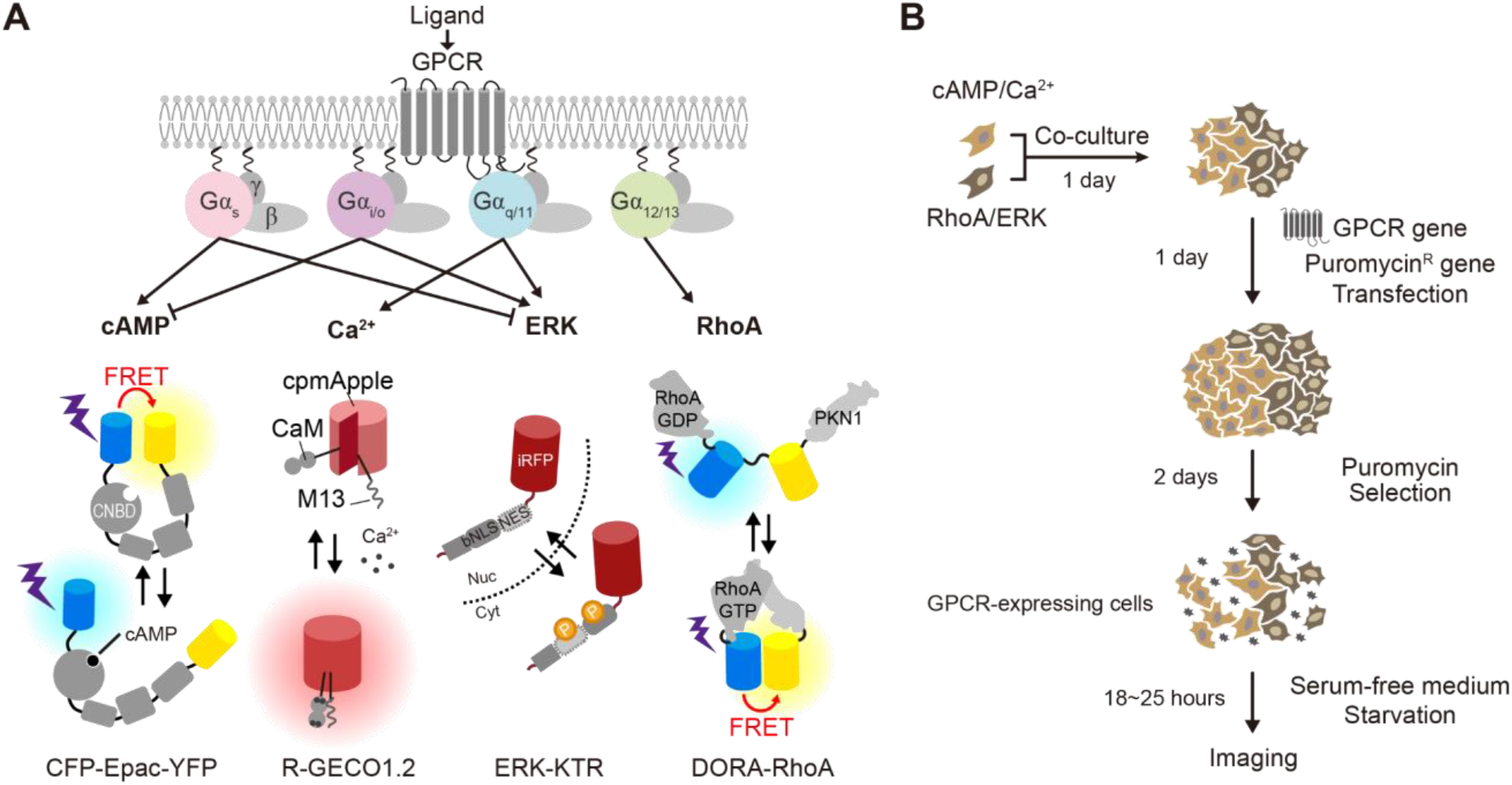
Development of the live-cell fluorescence imaging system for quantifying the GPCR signaling dynamics. (A) (Upper panel) Schematic model of Gα coupling and typical GPCR-mediated intracellular signaling pathways. (Lower panel) The mode of action of the indicated biosensors is shown. (B) Experimental procedure of the live-cell fluorescence imaging.

Much effort has been devoted to the development of imaging systems for quantifying GPCR activation upon ligand stimulation. The imaging systems reported so far can be roughly classified into two types: systems that utilize the principle of ligand-induced association/dissociation between GPCRs, effectors, and heterotrimeric G proteins, and systems that employ downstream signaling molecules of GPCRs as a marker. In the former imaging methods, protein-protein interactions such as GPCR-G proteins, GPCR-β-arrestin, and Gα-Gβγ are detected by techniques such as Förster (or fluorescence) resonance energy transfer (FRET) and bioluminescence resonance energy transfer (BRET) [15–17]. Other methods for measuring these protein-protein interactions by converting them into gene expression or by cleavage of membrane molecules by metalloproteinases such as ADAM17/TACE have been proposed to enable the large-scale GPCR screening [18,19]. These techniques directly evaluate ligand-induced GPCR activation, but they do not allow estimation of the extent to which the GPCR activation evokes downstream signaling. The latter type of imaging methods detect downstream cell signaling induced by GPCRs such as cAMP and Ca^2+^ by small chemical compounds and genetically encoded biosensors [20–23]. However, a drawback of this method is that it is challenging to visualize two or more types of signal transduction systems in an efficient manner, and the throughput is practically limited.

Many patterns of GPCR-Gα protein coupling have been investigated, revealing that some GPCRs can promiscuously couple to multiple G proteins [19,24]. Yet, in most cases, only a limited amount of quantitative information is available. Thus, it is still unknown to what extent the GPCRs activate Gα proteins and what types of downstream signaling dynamics an activated GPCR generates. This study reports a quantitative live-cell imaging system for quantifying cAMP, Ca^2+^, RhoA, and ERK activation dynamics induced by GPCR activation at the single-cell level. For this purpose, the FRET-based biosensors for cAMP and RhoA activity, the red fluorescent Ca^2+^ biosensor, and the infra-red fluorescent ERK biosensors were selected. We first established two types of HeLa cell lines stably expressing cAMP/Ca^2+^ biosensors and RhoA/ERK biosensors. These two cell lines were then co-cultured, transfected with GPCR genes, and stimulated with the ligand to monitor the changes in cAMP, Ca^2+^, RhoA, and ERK activity. Taking advantage of this method, we quantified the GPCR signaling dynamics of 5 dopamine receptors and 12 serotonin receptors, demonstrating a substantial difference even between GPCRs that bind to the same Gα proteins and cellular heterogeneity of downstream signals.

### Experimental

#### Plasmids

The cAMP biosensor (CFP-Epac-YFP) was developed based on the previous work [20], and it contained monomeric teal fluorescent protein (mTFP), the human RAPGEF3 (EPAC) gene (corresponding to amino acids 149-881) cloned from HeLa cells by RT-PCR, and mVenus. The cDNA of the cAMP biosensor was inserted into the pCX4neo vector [25], a retroviral vector with IRES-*neo* (the G418-resistance gene), providing pCX4neo-CFP-Epac-YFP. The cDNA of the red fluorescent genetically encoded Ca^2+^ indicators for optical imaging (R-GECO1.0) [26] was subcloned into the pPBbsr vector [27,28], a PiggyBac transposon vector with IRES-*bsr* (the blasticidin S-resistance gene), generating pPBbsr-R-GECO1.0. The dimerization-optimized reporter for activation (DORA) RhoA biosensor (DORA-RhoA) was used to visualize RhoA activity, and the DORA-RhoA was a gift from Dr. Yi Wu (University of Connecticut Health Center, CT) [22]; DORA-RhoA consists of the Rho-binding domain of PKN1, circularly permuted Venus (cpVenus), a linker, Cerulean3, and RhoA. The cDNA of DORA-RhoA was subcloned into pT2Absr [29], which was generated by inserting IRES-*bsr* into the pT2AL200R175 vector, generating pT2Absr-DORA-RhoA. The cDNA of the ERK activity biosensor, ERK-KTR [23], was fused with the cDNA of monomeric near-infrared fluorescent protein (miRFP703)[30], and subcloned into the pCSIIbleo vector, a lentivirus vector with IRES-*bleo* (the zeocin-resistance gene) [31], generating pCSIIbleo-ERK-KTR-miRFP703. The plasmid encoding the nuclear marker, Linker Histone-H1 (H1)-mCherry, was subcloned into a pCSIIneo vector [32], generating pCSIIneo-H1-mCherry. The GPCR plasmids were derived from the PRESTO-Tango kit originated from Dr. Bryan Roth (Addgene kit # 1000000068) [18]. pCMV-VSV-G-RSV-Rev was a gift of Dr. Miyoshi (RIKEN, Japan) [33], pGP and pCSIIpuro-MCS were a gift from Dr. Matsuda (Kyoto University, Japan) [34,35], psPAX2 was a gift from Dr. Trono (Addgene plasmid #12260) [36], pCAGGS-T2TP was a gift from Dr. Kawakami (National Institute for Genetics, Japan) [29], and pCMV-mPBase (neo-) was a gift from Dr. Bradley (Wellcome Trust Sanger Institute, UK) [27].

#### Reagents

Epidermal growth factor (EGF) was purchased from Sigma-Aldrich (St. Louis, MO). Blasticidin S, zeocin, G418, and puromycin were obtained from InvivoGen (Carlsbad, CA). Forskolin (FSK) and 3-isobutyl-1-methyl-xanthine (IBMX) were purchased from Wako (Osaka, Japan). Dopamine was obtained from Sigma-Aldrich (H8602) and solubilized into 10 mM HCl as a 1 M stock solution. Serotonin was purchased from Cayman (14332) and solubilized into 10 mM HCl as a 50 mM stock solution. Adenosine 5’-triphosphate disodium salt hydrate (ATP) was obtained from TCI (A0157) and solubilized into DDW as a 100 mM stock solution. Isoproterenol (ISO) was obtained from Sigma-Aldrich (I6504) and solubilized into DDW as a 50 mM stock solution.

#### Cell culture

HeLa cells and HEK-293T cells were gifted from Dr. Matsuda (Kyoto University, Japan), and cultured in Dulbecco’s Modified Eagle’s Medium (DMEM) high glucose (Wako; nacalai tesque) supplemented with 10% fetal bovine serum (Sigma-Aldrich) at 37°C in 5% CO_2_. For the live-cell imaging, HeLa cells were plated on CELLview cell culture dishes (glass bottom, 35 mm diameter, 4 components: The Greiner Bio-One) one day before transfection. Transfection was performed with a mixture containing 230 ng plasmid encoding TANGO-GPCR, 20 ng plasmid carrying the puromycin-resistance gene (pCSIIpuro-MCS), and 0.25 uL of 293fectin transfection reagent (Thermo Fisher Scientific) in each well. In the dopamine receptor co-expression experiment in Figure 4, TANGO GPCR plasmids were used in a 1: 1 mixture. One day after the transfection, puromycin (final concentration 2 μg/ml) was administered, and transfected cells were selected for 2 days. After the selection of transfected cells, the medium was replaced with starvation medium (FluoroBrite (nacalai tesque)/1x GlutaMAX (GIBCO)/0.1% BSA) 18 ~ 25 h before the imaging was started.

#### Stable cell line construction

To establish a stable cell line expressing R-GECO1.0 and CFP-Epac-YFP (HeLa/cAMP/Ca^2+^), the PiggyBac transposon system and the retroviral vector system were used, respectively. The pPBbsr-R-GECO1.0 was co-transfected with pCMV-mPBase (neo-) encoding the PiggyBac transposase by using 293fectin. One day after the transfection, cells were selected with 20 μg/ml blasticidin S for at least one week. For retroviral production, the pCX4neo-CFP-Epac-YFP was transfected into HEK-293T cells together with pGP and pCSV-VSV-G-RSV-Rev by using Polyethylenimine “MAX” MW 40,000 (Polyscience Inc., Warrington, PA). Virus-containing media were collected at 48 h after the transfection, filtered, and used to infect the cell lines expressing R-GECO1.0 with 10 μg/mL polybrene. After the infection, cells were selected with 20 μg/ml blasticidin S and 1 mg/mL G418 for at least one week, followed by single-cell cloning. Clonal cells were used in the experiments Figure 2 and Figure 4.

**Figure 2.**
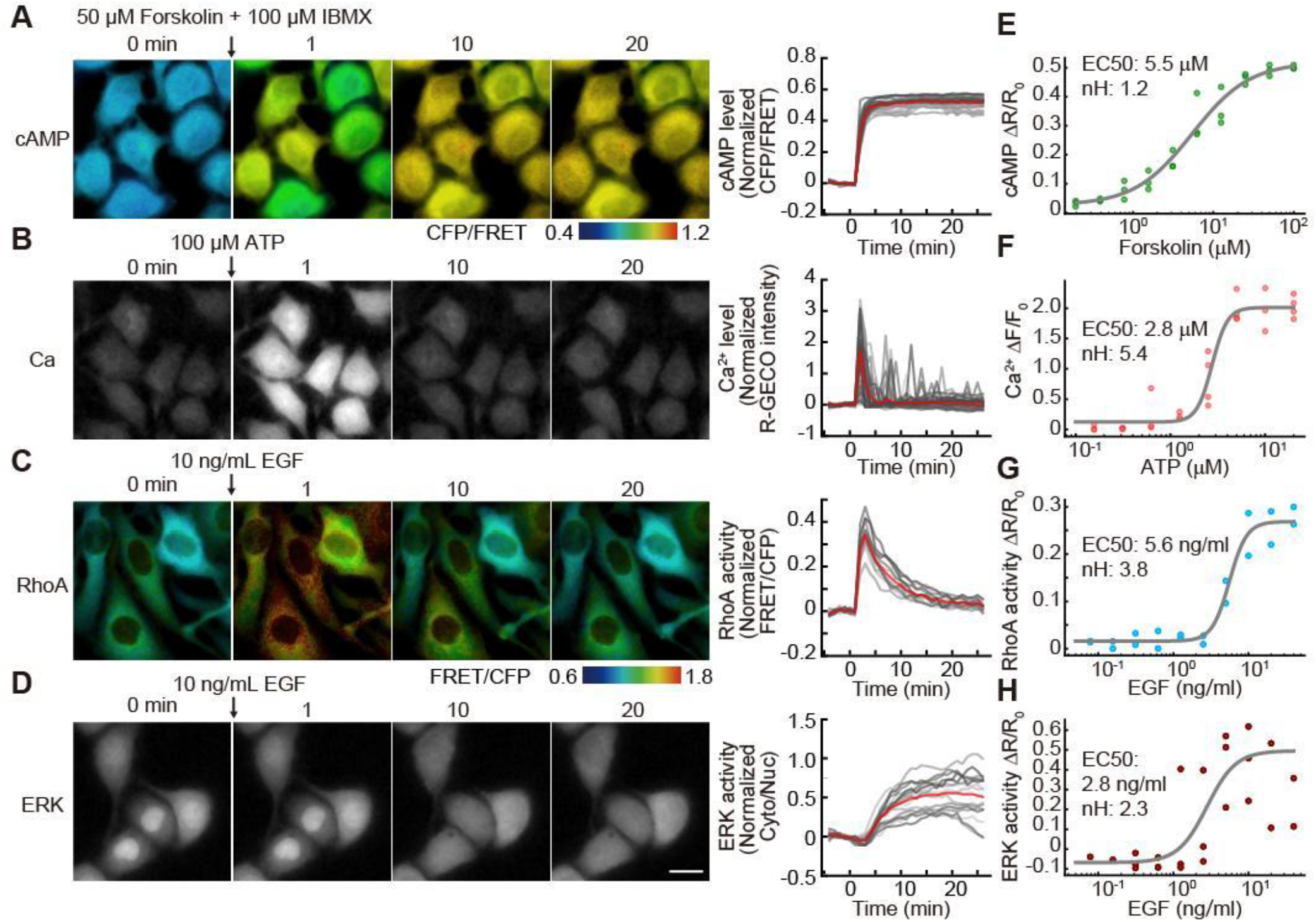
Validation of cell lines for monitoring GPCR signaling. (A-D) (Left panels) HeLa/cAMP/Ca^2+^ cells (A, B) and HeLa/RhoA/ERK cells (C, D) were treated with 50 μM Forskolin and 100 μM IBMX (A), 100 μM ATP (B), and 10 ng/mL EGF (C and D). Representative CFP/FRET ratio images (A) and FRET/CFP ratio images (C) are shown in the IMD mode, where eight colors from red to blue are used to represent the CFP/FRET ratio (A) or FRET/CFP ratio (C) with the intensity of each color indicating the denominator channels. (Right panels) The response of the biosensors for cAMP levels, Ca^2+^ levels, ERK activity, and RhoA activity is normalized by dividing by the averaged value before stimulation and plotted as a function of elapsed time after stimulation. The red and gray lines represent the time-course for the average and individual cells, respectively. N = 32, 45, 13, 16 cells, respectively. (E-H) The dose response curve of cAMP (E), Ca^2+^ (F), RhoA (G), and ERK (H) to the indicated ligands. The mean ΔF/F_0_ or ΔR/R_0_ values (dots) are plotted as a function of ligand concentrations with the fitted curve (gray line) and parameters for EC50 and Hill coefficient (nH). The experiments were repeated at least two times. Scale bar, 10 μm.

To establish a stable cell line expressing ERK-KTR-miRFP703, DORA-RhoA, and H1-mCherry (HeLa/RhoA/ERK), a lentiviral vector system (for ERK-KTR-miRFP703 and H1-mCherry) and a Tol2 transposon-based system (for DORA-RhoA) were applied, respectively. For lentiviral production, HEK-293T cells were co-transfected with the pCSIIbleo-ERK-KTR-miRFP703, psPAX2, and pCSV-VSV-G-RSV-Rev by using Polyethylenimine “MAX” MW 40,000. Virus-containing media were collected at 48 h after the transfection, filtered, and used to infect HeLa cells with 10 μg/mL polybrene. After the infection, cells were selected with 100 μg/ml zeocin for at least one week. Next, the pT2Absr-DORA-RhoA was co-transfected into HeLa cells stably expressing ERK-KTR-miRFP703 together with the pCAGGS-T2TP plasmid encoding transposase by using 293fectin. One day after the transfection, cells were selected by at least one week of treatment with 100 μg/ml zeocin and 20 μg/ml blasticidin S. The pCSIIneo-H1-mCherry was introduced into this cell line by using a lentiviral viral vector system as well as ERK-KTR-miRFP703. One day after the infection, cells were selected by at least one week of treatment with 100 μg/ml zeocin, 20 μg/ml blasticidin S, and 1 mg/mL G418, followed by single-cell cloning.

Single-cell clones were used in the experiments in Figure 2, Supplementary Figures S2, S3, and S5. In other experiments, bulk cells were used.

#### Time-lapse imaging

Images were acquired on an IX81 inverted microscope (Olympus) equipped with a Retiga 4000R cooled Mono CCD camera (QImaging), a Spectra-X light engine illumination system (Lumencor), an IX2-ZDC laser-based autofocusing system (Olympus), a UAPO/340 40x/1.35 oil iris objective lens (Olympus), a MAC5000 controller for filter wheels and XY stage (Ludl Electronic Products), an incubation chamber (Tokai Hit), and a GM-4000 CO_2_ supplier (Tokai Hit). The following filters and dichroic mirrors were used: for R-GECO1.0 and mCherry, an FF01-580/20 excitation filter (Semrock), a 20/80 beamsplitter dichroic mirror (Chroma), and an FF01-641/75 emission filter (Semrock); for iRFP, an FF01-632/22 excitation filter (Semrock), an FF408/504/581/667/762-Di01 dichroic mirror (Semrock), and an FF01-692/LP emission filter (Semrock); for FRET, an FF01-438/24 excitation filter (Semrock), an XF2034 455DRLP dichroic mirror (Omega Optical), an FF01-542/27 emission filter (Semrock); for CFP, an FF01-438/24 excitation filter (Semrock), an XF2034 455DRLP dichroic mirror (Omega Optical), and an FF01-483/32 emission filter (Semrock). The microscopes were controlled by MetaMorph software (Molecular Devices). For live-cell imaging, cells were stimulated with ligands (EGF, FSK+IBMX, ATP, dopamine, or serotonin) 5 min after the start of imaging, and followed by 25 min imaging with every 1 min interval. To measure the Gα_i/o_ activity, cells were first treated with 100 nM ISO 5 min after the start of imaging to increase the intracellular cAMP level. Next, cells were subsequently treated with vehicle or ligands (dopamine or serotonin) 10 min after the ISO treatment, followed by 15 min imaging.

#### Immunofluorescence

Cells were fixed with the final 3.7% formaldehyde for 30 min in a culture medium. After washing with PBS four times, cells were permeabilized with 0.2% TritonX-100 in PBS for 15 min for the immunofluorescence of total GPCR expression, followed by soaked with 3% BSA and 0.02% TritonX-100 in PBS (buffer 1). For the immunofluorescence of GPCRs expressing cell-surface, cells were soaked with buffer 1 without permeabilization. Then, the cells were incubated with primary antibodies, monoclonal anti-FLAG M2 antibody (1:500 dilution; Sigma-Aldrich, F1804) diluted in buffer 1 for overnight at 4°C. Next, the cells were washed three times with PBS, and then incubated for 1 h at room temperature with Goat anti-Mouse IgG (H+L) Highly Cross-Adsorbed Secondary Antibody Alexa Fluor Plus 555 (1:1000 dilution; Invitrogen, A32727) in buffer 1. Finally, the cells were washed three times with PBS and subjected to fluorescence imaging.

#### Image and data analysis

Fiji, a distribution of ImageJ [37], was used for image processing and analysis. For all images, background signals were subtracted by the rolling ball method, and then the images were registered by StackReg, a Fiji plugin to correct misregistration. Note that the median filter was used for the time-lapse images before registration to remove camera noise to prevent the registration error. For FRET images of CFP-Epac-YFP or DORA-RhoA, the ratio images of the CFP/FRET or FRET/CFP ratio were generated, respectively, and then ratio images were visualized by the intensity-modulated display (IMD) mode, where eight colors from red to blue are used to represent the CFP/FRET ratio.

To measure the intracellular cAMP and Ca^2+^ levels, we manually set a circular region of interest (ROI) with a diameter of 20 pixels at the center of each cell. Note that we avoid selecting the cells that overlapped with dying cells during the time course. To measure the ERK activity from the translocation of ERK-KTR, we used CellTK [38]. In brief, pre-processing, segmentation, tracking, and post-processing was performed using functions in CellTK. In pre-processing, nuclear images were processed by a gaussian blur filter. In the segmentation process, an adaptive threshold algorithm was basically applied except for some difficult images, which were segmented by Stardist [39,40]. Segmented nuclear labels were tracked by the Linear Assignment Problem (LAP) algorithm [41], followed by a track_neck_cut function in CellTK. As for post-processing, short tracking data were discarded, and the cytoplasmic region was defined as a dilating ring around nuclear segmentation. Quantified feature values (i.e., mean intensity, x-y positions, etc.) of each cell were exported and cleaned in Python. Invalid cell tracks were discarded according to the following criteria: a threshold of CFP signal intensity to remove non-fluorescent cells and dead cells; a threshold of coefficient of variation (CV) of the CFP signal intensity along time course to remove the cells crossed by floating dead cells or debris during time course; DORA-RhoA cells adjacent to CFP-Epac-YFP cells were removed because the much stronger fluorescence from CFP-Epac-YFP cells affects the quantification of FRET signals of DORA-RhoA. The quantified time traces were further normalized by the average values before stimulation (0-5 min), except in Figure 6E and Figure 7F.

To classify signaling dynamics, *K*-means-based time-series clustering was used. Euclidean distance was used as a measure of proximity. At first, all quantified time-course data of each reporter were normalized as described above, and divided into three groups by the *K*-means algorithm using scikit-learn ver. 1.0.2 and tslearn ver. 0.5.2 [42,43] modules for Python. At the next step, the fractions of clusters were calculated in each receptor. Note that the number of resulting clusters was conservatively determined as three for all reporters because the cluster number prediction methods, such as the Elbow Method, did not provide reliable prediction in our case.

## RESULTS

### Development of live-cell fluorescence imaging for the quantification of GPCR signaling

To cover a wide range of GPCR signals mediated by individual Gα subtypes, we chose cAMP, Ca^2+^, RhoA, and ERK as markers (**Figure 1A**). We screened genetically encoded biosensors for these signaling molecules, and finally selected the following biosensors: CFP-Epac-YFP (a Förster resonance energy transfer (FRET)-based cAMP biosensor) [20], R-GECO1.0 (a red fluorescent Ca^2+^ biosensor) [21], DORA-RhoA (a FRET-based RhoA biosensor) [22], and ERK-KTR fused with miRFP703 (an infra-red fluorescent ERK biosensor) [23] (**Figure 1A**). For efficient imaging, two types of stable HeLa cell lines were established: a cell line expressing the cAMP biosensor and R-GECO1.0 (HeLa/cAMP/Ca^2+^), and a cell line expressing DORA-RhoA and ERK-KTR (HeLa/RhoA/ERK) (**Figure 1B**) together with linker histone H1-mCherry as a nuclear marker to quantify the cytoplasm/nucleus (C/N) ratio automatically. The two cell lines were co-cultured and transfected with the plasmids for GPCR expression and puromycin-resistance gene expression. One day after transfection, the cells were treated with puromycin to select GPCR-expressing cells. After the selection, the cells were further serum-starved for 18~25 h before imaging to reduce the ERK activity at a basal level. The plasmids from the PRESTO-Tango GPCR kit were used for the expression of each GPCR, and therefore the exogenous GPCR protein was fused with the HA signal peptide and FLAG-tag at the N-terminus and V2 tail, the TEV cleaved site, and the rtTA moieties at the C-terminus [18]. We suspect that these parts have no considerable effects on GPCR signaling.

### Validation of cell lines for monitoring GPCR signaling

As a validation of the cell lines mentioned above, we stimulated the cells with chemical compounds, ligands, and epidermal growth factor (EGF), which are well-known to upregulate cAMP, Ca^2+^, RhoA, and ERK. HeLa/cAMP/Ca^2+^ cells were stimulated with Forskolin and IBMX (**Figure 2A**) and ATP (**Figure 2B**). The cells showed a sustained increase in the CFP/FRET ratio of the CFP-Epac-YFP biosensor and pulsatile increase in red fluorescence of R-GECO1.0, indicating a sustained cAMP increase and pulsatile Ca^2+^ increase, respectively. Similarly, HeLa/RhoA/ERK cells were stimulated with EGF, resulting in a transient rise in the FRET/CFP ratio of DORA-RhoA and gradual translocation of ERK-KTR-miRFP703 from the nucleus to the cytoplasm (**Figure 2C, 2D**). The data indicate that EGF induced transient RhoA and gradual ERK activation. We represent ERK activity as the cytoplasm/nucleus (C/N) ratio of ERK-KTR-miRFP703 fluorescence intensity [23,30]. To assess the sensitivity and dynamic range of biosensors, we evaluated the dose-response of each biosensor to the ligands (**Figure 2E-2H**). Further, no clear correlation was observed between the expression level of the biosensors and the response to the ligands in the cell lines we used in this study (**Supplementary Figure S1A-D**). It is noted that our microscope setting did not show any spectral cross-talk between CFP-Epac-YFP and R-GECO1.0 or between DORA-RhoA and ERK-KTR-miRFP703. Taken together, these results demonstrated that the cell lines expressing biosensors allowed monitoring of cAMP, Ca^2+^, RhoA, and ERK activity at the single-cell level.

### Signaling dynamics of dopamine receptors

A neuromodulator, dopamine, controls many physiological functions, including locomotion, reward, cognitive function, and learning [44]. Because dopamine is involved in a wide range of cellular and behavioral processes, multiple human disorders have been linked to dopaminergic dysfunctions such as Parkinson’s disease, schizophrenia, ADHD, and Tourette’s syndrome [45]. The five subtypes of dopamine receptors, D1, D2, D3, D4, and D5 receptors, are encoded by the genes *DRD1*, *DRD2*, *DRD3*, *DRD4*, and *DRD5*, respectively, in the human genome, and collectively these subtypes are known to mediate all physiological functions of dopamine [45,46]. The dopamine receptors are GPCRs and are divided into two groups: the D1-class dopamine receptors (D1 and D5) and the D2-class dopamine receptors (D2, D3, and D4) [47]. The D1-class dopamine receptors are generally coupled to Gα_s_, while the D2-class dopamine receptors are coupled to Gα_i/o_ (**Figure 3A**). Thus, the D1 and D2 receptors positively and negatively regulate adenylate cyclases (ACs), which catalyze the production of cAMP, respectively. Furthermore, it has been reported that D1, D5, and a putative D1-D2 heterodimer couple and activate Gα_q/11_ and Ca^2+^ signaling [48–51]. However, this idea remains controversial because D1and D2 are not coexpressed in most striatal neurons in mice [45], and further studies will be needed to quantitatively investigate to extent to which these dopamine receptors contribute to the Gα_q/11_ activation and Ca^2+^ increase.

**Figure 3.**
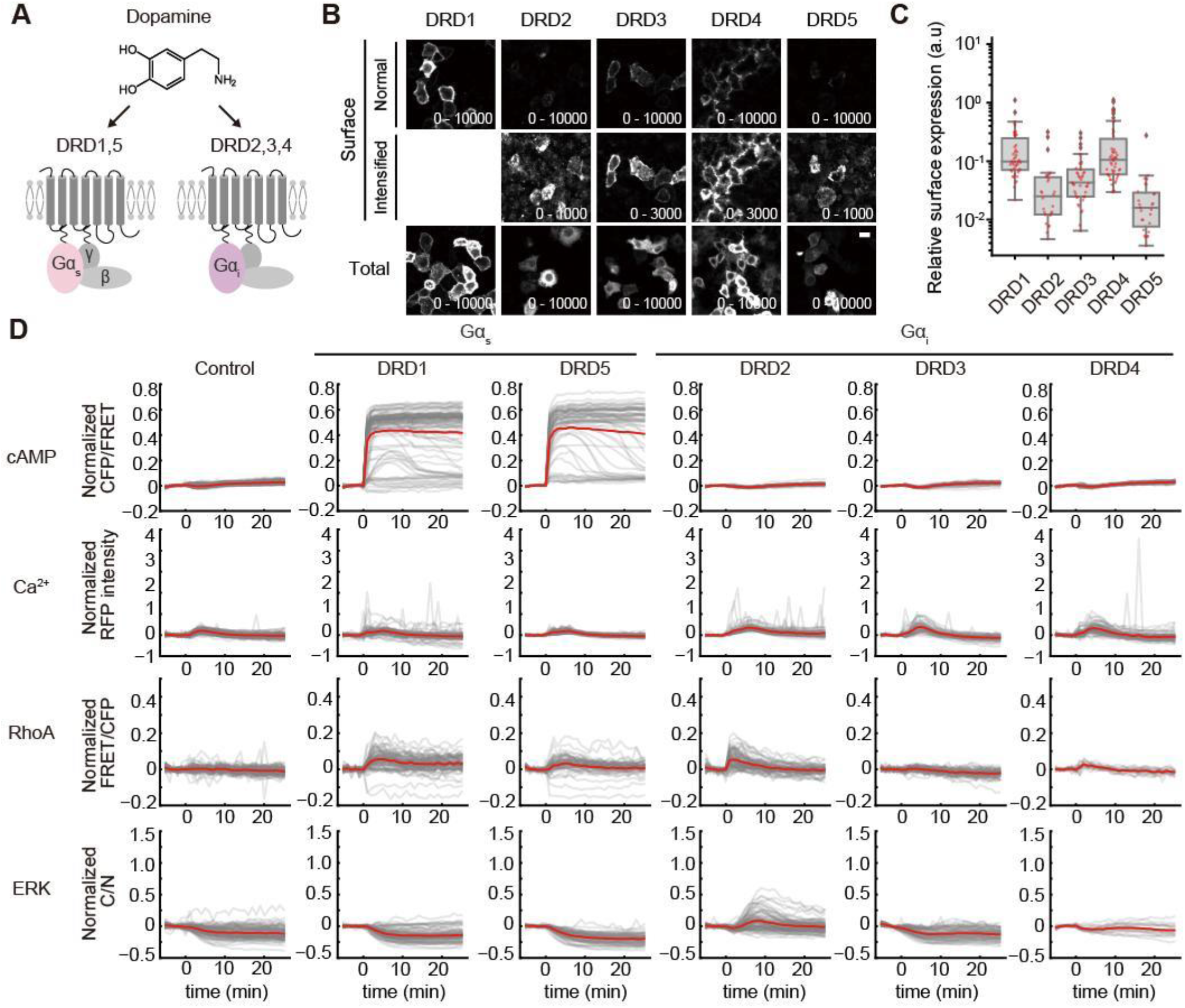
Signaling dynamics induced by each dopamine receptor. (A) Schematic illustration of dopamine, dopamine receptors (DRDs) coupling to Gα, and downstream signaling. (B) Expression of DRDs at the cell surface and whole cell. HeLa cells were transfected with the plasmids expressing the indicated receptors, followed by immunofluorescence (see methods). The values in the lower right indicate the lower and upper limits of the display range. Scale bar, 20 μm. (C) The quantification of the surface expression levels of DRDs. To correct for cell-to-cell heterogeneity of transfection efficiency, the fluorescence intensity of cotransfected EGFP was used to normalize the fluorescence intensity of cell surface receptors in each cell. The values for each cell were plotted as a scatter plot with a box-whisker plot, in which the box shows the quartiles of data with the whiskers denoting the minimum and maximum except for the outliers detected by 1.5 times the interquartile range (n = 46, 26, 41, 50, 27 cells). (D) Normalized responses of each reporter in HeLa cells expressing the indicated dopamine receptors are shown. The response of the biosensors for cAMP levels, Ca^2+^ levels, ERK activity, and RhoA activity is normalized by dividing by the averaged value before stimulation and plotted as a function of elapsed time after administration of 10 μM dopamine. The red and gray lines represent the time-course for the average and individual cells (n > 10 cells), respectively.

First, the surface expression of D1-5 receptors was confirmed by immunofluorescence assay (**Figure 3B and 3C**). DRD1 was mainly localized at the cell surface, while DRD3 and DRD4 were partially expressed in the cell surface and the rest was localized intracellularly (**Figure 3B**). Cell surface expression of DRD2 and DRD5 were weaker, but detectable, than the others, and most of these receptors retained in the intracellular regions (**Figure 3B**). We quantified the surface expression levels of DRDs by correcting cell-to-cell heterogeneity of transfection efficiency, showing the distinct differences of surface expression levels among the DRDs (**Figure 3C**). Next, we quantified the cAMP, Ca^2+^, RhoA, and ERK activation dynamics evoked by dopamine receptors with the live-cell fluorescence imaging system (**Figure 1B**). In HeLa cells lacking expression of dopamine receptors, dopamine stimulation did not elicit any change in cAMP, Ca^2+^, RhoA, or ERK activity (**Figure 3D**, left column). This could be simply because HeLa cells do not express any dopamine receptors. As expected, the expression of D1-type receptors, i.e., DRD1 and DRD5, increased cAMP levels upon dopamine stimulation, while the expression of D2-type receptors, i.e., DRD2, DRD3, and DRD4, did not change any cAMP levels (**Figure 3D**, first row). This was probably because the basal cAMP level was low, and thus the cAMP biosensor could not detect the decrease in cAMP levels by D2-type receptors. Weak, measurable Ca^2+^ pulses were observed in HeLa cells expressing DRD1 and DRD2, suggesting that DRD1 and DRD2 receptors are weakly coupled to Gα_q/11_ receptors (**Figure 3D**, second row). RhoA was also activated in DRD1 or DRD5 expressing HeLa cells for an unknown reason (**Figure 3D**, third row). ERK was transiently decreased in HeLa cells expressing DRD1 or DRD5, but it is challenging to evaluate dopamine-induced ERK signaling due to significant cell heterogeneity (**Figure 3D**, fourth row). Furthermore, to confirm the activation of Gα_i/o_-coupled receptors, cells expressing DRD2, DRD3, or DRD4 were pretreated with 100 nM isoproterenol to upregulate cAMP production through endogenous β adrenoceptor activation, followed by dopamine stimulation. As we expected, the expression of DRD2, 3, or 4 increased the proportion of cells that reduced cAMP levels after dopamine stimulation (**Supplementary Figure S2A**), indicating the dopamine-induced activation of Gα_i/o_ signaling through DRD2, 3, and 4.

Next, we examined whether heterodimerized dopamine receptors synergistically increase intracellular Ca^2+^ levels upon dopamine stimulation, because it has been reported that heterodimers of dopamine receptors such as the D1-D2 dimer activate Gα_q/11_ signaling [48–50]. HeLa cells were co-transfected with ten possible combinations of dopamine receptors and stimulated with dopamine. Unexpectedly, we found no enhancement of the dopamine-induced Ca^2+^ increase by co-expression of dopamine receptors, but rather the co-expression inhibited the pulsatile Ca^2+^ increase observed in HeLa cells expressing DRD1 or DRD5 (**Supplementary Figure S3**).

### Signaling dynamics of serotonin receptors

Serotonin (5-hydroxytryptamine, 5-HT) plays essential roles in a wide range of physiological phenomena, including sensory perception and behaviors [52]. The serotonin receptors are the most prominent among the GPCR families for neurotransmitters; 13 distinct genes encode GPCRs for the serotonin, and there is one ligand-gated ion channel, the 5-HT3 receptor [53]. The G-protein coupled serotonin receptors are grouped into three families depending on which Gα proteins they primarily couple: the Gα_q/11_-coupled receptors (HTR2A, HTR2B, and HTR2C), Gα_i/o_-coupled receptors (HTR1A, HTR1B, HTR1D, HTR1E, HTR1F, HTR5), and Gα_s_-coupled receptors (HTR4, HTR6, and HTR7) (**Figure 5A**). Although these receptors are divided based primarily on their coupling with Gα proteins, recent works have shown that GPCRs can couple to more than one type of Gα proteins [19]. In addition, the constitutive activity of serotonin receptors has been reported, though the basal- and ligand-induced activities of serotonin receptors have not yet been thoroughly evaluated.

We therefore analyzed the G-protein-coupled serotonin receptor-induced cAMP, Ca^2+^, RhoA, and ERK activation with the live-cell fluorescence imaging system (**Figure 1B**). First, Gα_q/11_-coupled serotonin receptors of HTR2A, HTR2B, and HTR2C, were investigated as did in DRDs. The surface and total expression of HTR2A and HTR2B were detectable, while those of HTR2C were quite low (**Figure 4B and 4C**). Parental HeLa cells did not show any response of these signaling molecules to serotonin stimulation (**Figure 4D**, left column). The expression of Gα_q/11_-coupled HTR2A and HTR2C induced a sustained or pulsatile Ca^2+^ increase upon 5-HT stimulation, while slight or no response of Ca^2+^ was observed in HTR2B-expressing cells (**Figure 4D**, second row). 5-HT stimulation slightly activated RhoA in HTR2A- or HTR2B-expressing cells (**Figure 4D**, third row). Some cells showed a substantial change in ERK activity, but on average, many cells did not show a shift in ERK activity (**Figure 4D**, fourth row).

**Figure 4.**
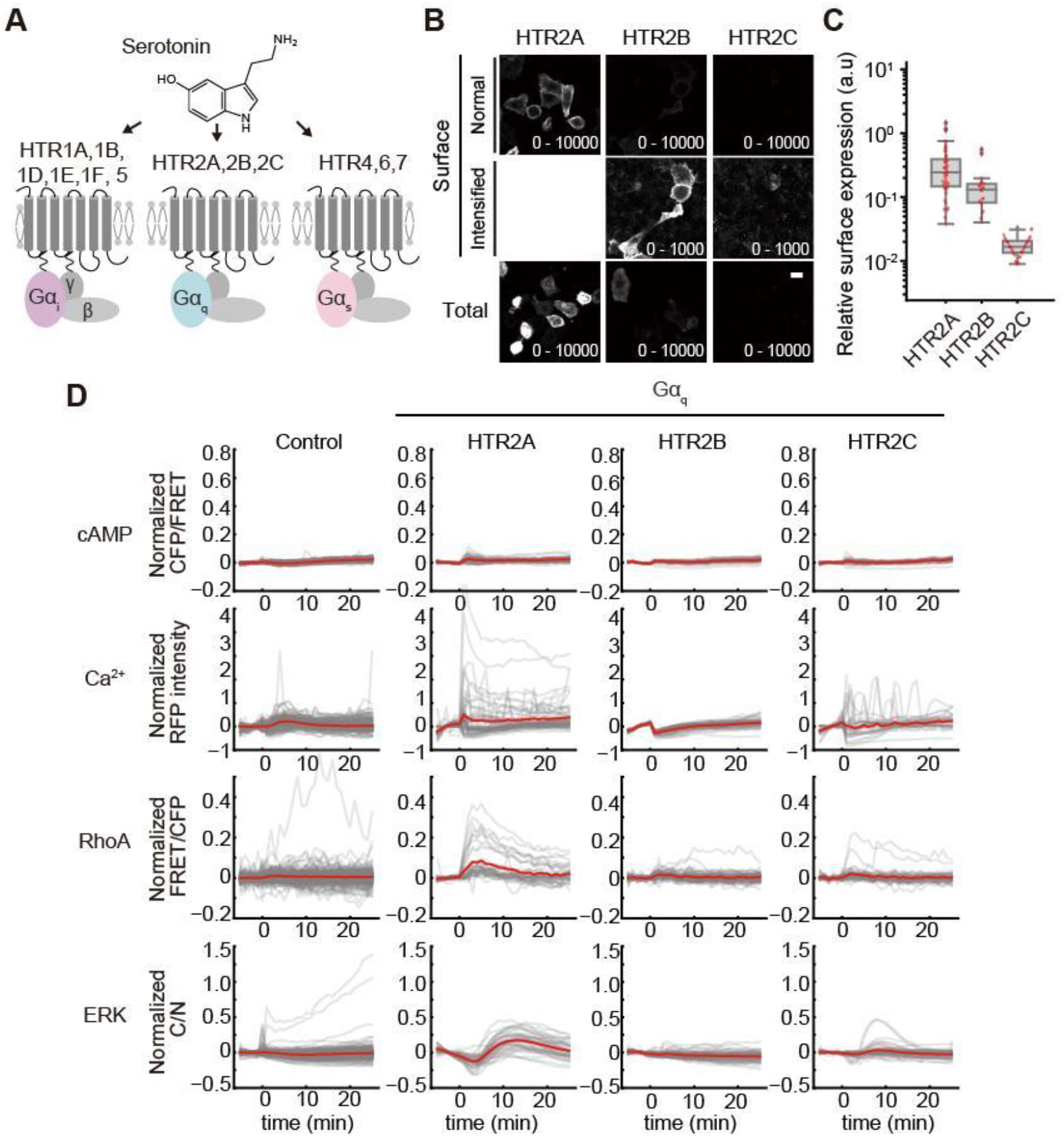
Signaling dynamics of Gα_q/11_-coupled serotonin receptors. (A) Schematic illustration of serotonin receptors coupling to Gα and downstream signaling. (B) Expression of Gα_q/11_-coupled serotonin receptors at the cell surface and whole cell. The data were obtained and are represented as in Figure 3B. Scale bar, 20 μm. (C) The quantification of the surface expression levels of Gα_q_-coupled serotonin receptors. The data were quantified and are represented as in Figure 3C. (n = 50, 21, 32 cells). (D) Normalized response of each reporter in HeLa cells overexpressing Gα_q/11_-coupled serotonin receptors was plotted as a function of time after 10 μM 5-HT administration. The response of the biosensors for cAMP levels, Ca^2+^ levels, ERK activity, and RhoA activity was normalized by dividing by the averaged value before stimulation and plotted as a function of elapsed time after stimulation. The red and gray lines represent the time-course for the average and individual cells (n > 10 cells), respectively.

We next analyzed the signaling dynamics of Gα_i/o_-coupled serotonin receptors, i.e., HTR1A, HTR1B, HTR1D, HTR1E, HTR1F, and HTR5. The surface and total expression of these receptors were confirmed by immunofluorescence (**Figure 5A and 5B**). As in D2-type dopamine receptors, the expression of these receptors did not alter the cAMP levels with or without 5-HT (**Figure 5C**, first row). Some of the receptors induced an increase in the Ca^2+^ level (HTR1A, 1B, 1D)(**Figure 5C**, second row) and RhoA activity (HTR1B)(**Figure 5C**, third row), indicating the possibility of Gα_q/11_-coupling signaling in these receptors. We could not observe the characteristic dynamics of ERK activation with high reproducibility (**Figure 5C**, fourth row). To confirm the activation of Gα_i/o_, isoproterenol was pretreated with the cells expressing the Gα_i/o_-coupled serotonin receptors, followed by the treatment of 5-HT. To our surprise, we found that a part of the HeLa cell population responded to 5-HT without overexpression of Gα_i/o_-coupled serotonin receptors (**Supplementary Figure 2B**). This could be due to the expression of HTR1D in HeLa cells [54]. The expression of the Gα_i/o_-coupled serotonin receptors further increased the population of responded cells to 5-HT except for HTR1E-expressing cells (**Supplementary Figure 2B**).

**Figure 5.**
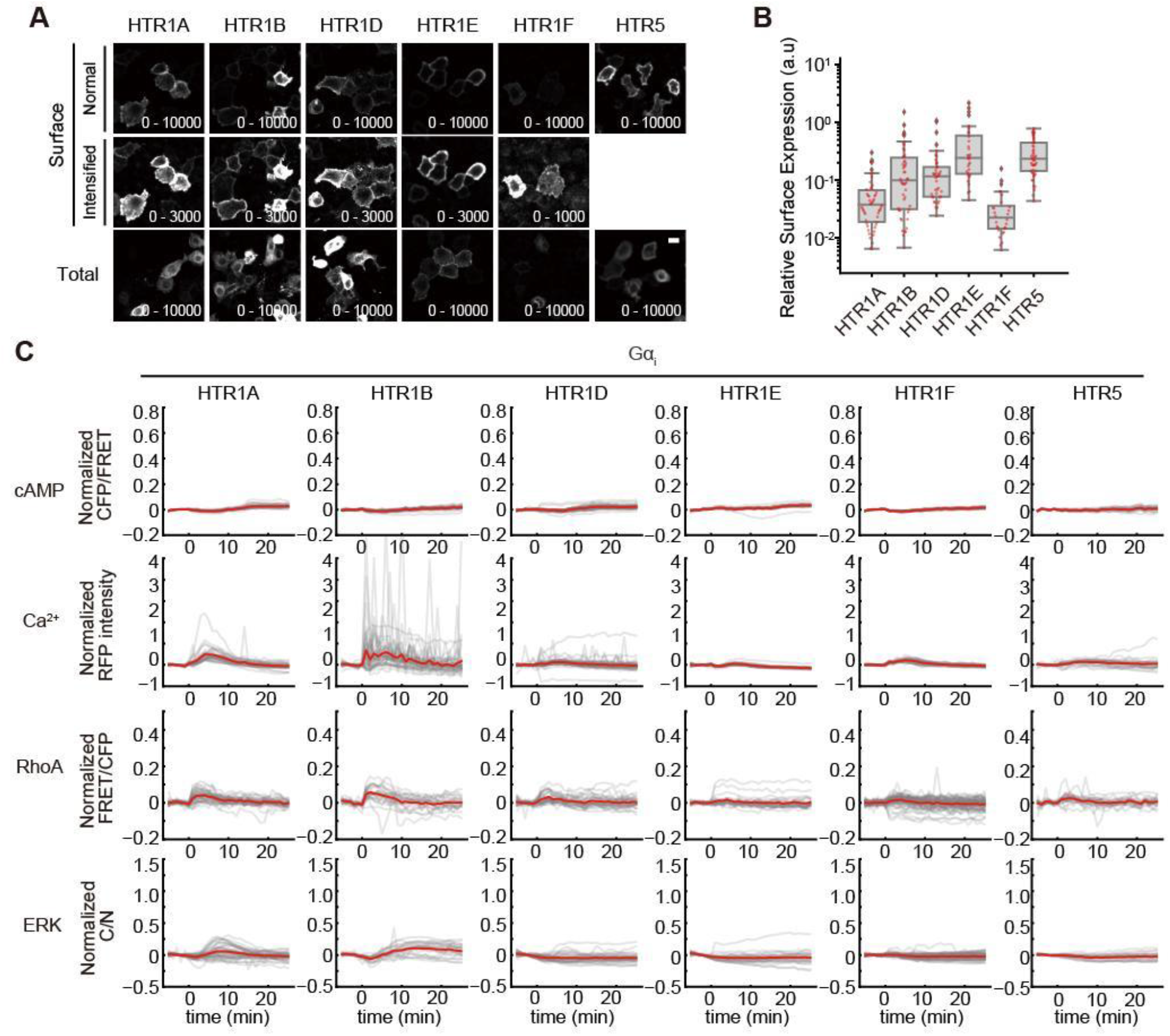
Signaling dynamics of Gαi/o-coupled serotonin receptors. (A) Expression of Gα_i/o1_-coupled serotonin receptors at the cell surface and whole cell. The data were obtained and are represented as in Figure 3B. Scale bar, 20 μm. (B) The quantification of the surface expression levels of Gα_i/o_-coupled serotonin receptors. The data were quantified and are represented as in Figure 3C. (n = 50, 50, 41, 32, 28, 50 cells). (C) Normalized response of each reporter in HeLa cells overexpressing Gα_i/o_-coupled serotonin receptors was plotted as a function of time after 10 μM 5-HT administration. The response of the biosensors for cAMP levels, Ca^2+^ levels, ERK activity, and RhoA activity was normalized by dividing by the averaged value before stimulation and plotted as a function of elapsed time after stimulation. The red and gray lines represent the time-course for the average and individual cells (n > 10 cells), respectively. (E) FSK → Ligand

### Constitutive activity of Gαs-coupled serotonin receptors and their response to ligand

Finally, we analyzed Gα_s_-coupled serotonin receptors, HTR4, HTR6, and HTR7. These receptors were expressed well at the cell surface and whole-cell (**Figure 6A and 6B**). HeLa cells expressing HTR4, HTR6, or HTR7 showed a sustained increase in cAMP level upon 5-HT stimulation with substantial cell-to-cell heterogeneity (**Figure 6C**, first row). In most cells, Ca^2+^ levels did not vary by 5-HT treatment (**Figure 6C**, second row), whereas RhoA activity slightly but reproducibly increased in cells expressing HTR4, HTR6, or HTR7 (**Figure 6C**, third row). HTR4-expressing cells showed an increase in ERK activity from the basal level on 5-HT stimulation, while did not in HTR6- or HTR7-expressing cells (**Figure 6C**, fourth row).

**Figure 6.**
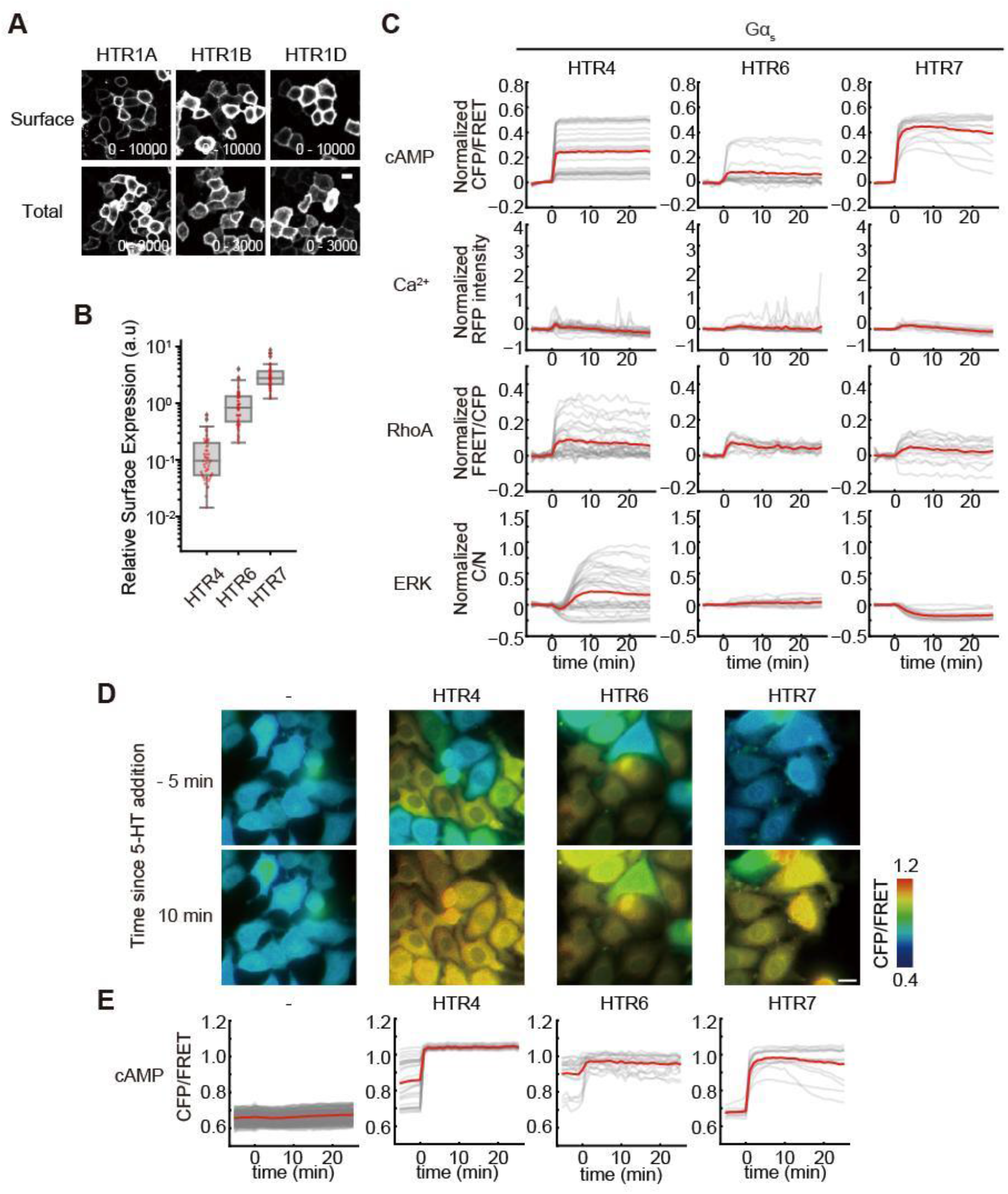
Constitutive activity of Gα_s_-coupled serotonin receptors and their response to ligand. (A) Expression of Gα_s_-coupled serotonin receptors at the cell surface and whole cell. The data were obtained and are represented as in Figure 3B. Scale bar, 20 μm. (B) The quantification of the surface expression levels of Gα_i/o_-coupled serotonin receptors. The data were quantified and are represented as in Figure 3C. (n = 50, 50, 50 cells). (C) Normalized response of each reporter in HeLa cells overexpressing Gα_s_-coupled serotonin receptors was plotted as a function of time after 10 μM 5-HT administration. Gray lines indicate single-cell responses (n > 10 cells), and the red line represents the mean response. (D) Representative CFP/YFP ratio images of cAMP in HeLa cells overexpressing Gα_s_-coupled serotonin receptors before and after the stimulation of 10 μM 5-HT. CFP/FRET ratio images are shown in intensity-modulated display mode as in Figure 2. Scale bar 10 μm. (E) Time course of CFP/FRET ratio values without normalization. Gray lines indicate single cell (n > 10 cells) responses, and the red line represents the mean response.

We investigated the cause of the large cell-to-cell heterogeneity in the increase in cAMP, and found that the basal cAMP levels varied between cells. HTR4 or HTR6 expression enhanced the basal cAMP level in some cells despite the absence of 5-HT stimulation (**Figure 6D**, upper panels). The cells with lower basal cAMP levels increased the cAMP upon 5-HT stimulation, whereas the cells with higher basal cAMP levels did not (**Figure 6D**, lower panels). As a result, 5-HT stimulation caused constant cAMP levels in all cells (**Figure 6D**, lower panels). Raw values of the CFP/YFP ratio of the cAMP biosensor, which correlated with intracellular cAMP concentration, also demonstrated that 5-HT stimulation kept cAMP levels constant (**Figure 6E**). These results suggest the constitutive activity of HTR4 and HTR6 for Gα_s_ signaling in the basal state and homeostatic mechanisms that control cAMP levels in the activating state.

### Systems analysis of signaling dynamics induced by activating dopamine or serotonin receptors

To take full advantage, we analyzed the time-series data of GPCR signaling in more detail. We performed hierarchical clustering with all data and demonstrated three clusters in the single-cell imaging data of cAMP, Ca^2+^, RhoA, and ERK (**Figure 7A-D**, left heatmaps and middle plots). Next, we quantified the fraction of these three clusters in the signaling dynamics of each GPCR (**Figure 7A-7D**, **Supplementary Figure S4A-D**). All Gα_s_-coupled receptors including DRD1, DRD5, HTR4, HTR6, and HTR7 demonstrated distinct cAMP dynamics in comparison to the other Gα-coupled receptors, though the fraction of the clusters in the cAMP dynamics varied among these Gα_s_-coupled receptors (**Figure 7A**, **Supplementary Figure S4A**). Some cells showing characteristic Ca^2+^ dynamics were found in DRD2, HTR1A, HTR1B, HTR2A, and HTR2C coupled to Gα_i/o_ or Gα_q/11_ (**Figure 7B**, **Supplementary Figure S4B**). Many dopamine or serotonin receptors affected the activity of RhoA, and among them, HTR2A, HTR4 and HTR7 demonstrated higher proportion of cells in cluster 3 showing strong activation of RhoA (**Figure 7C**, **Supplementary Figure S4C**). Interestingly, a fraction of cells changing RhoA activity tended to be higher in activation of Gα_s_-coupled GPCRs than the other Gα-coupled GPCRs. Although in control cells the proportion of cluster 2 showing ERK inactivation was observed, the distinct patterns of clustering in ERK activity dynamics were induced by the activation of DRD1, DRD5, HTR4, HTR7, and HTR2A (**Figure 7D**, **Supplementary Figure S4D**). These results suggest that GPCRs induce various downstream signaling dynamics, and that they contain some parts that can be explained by the Gα protein to which they are coupled and some that are not.

**Figure 7.**
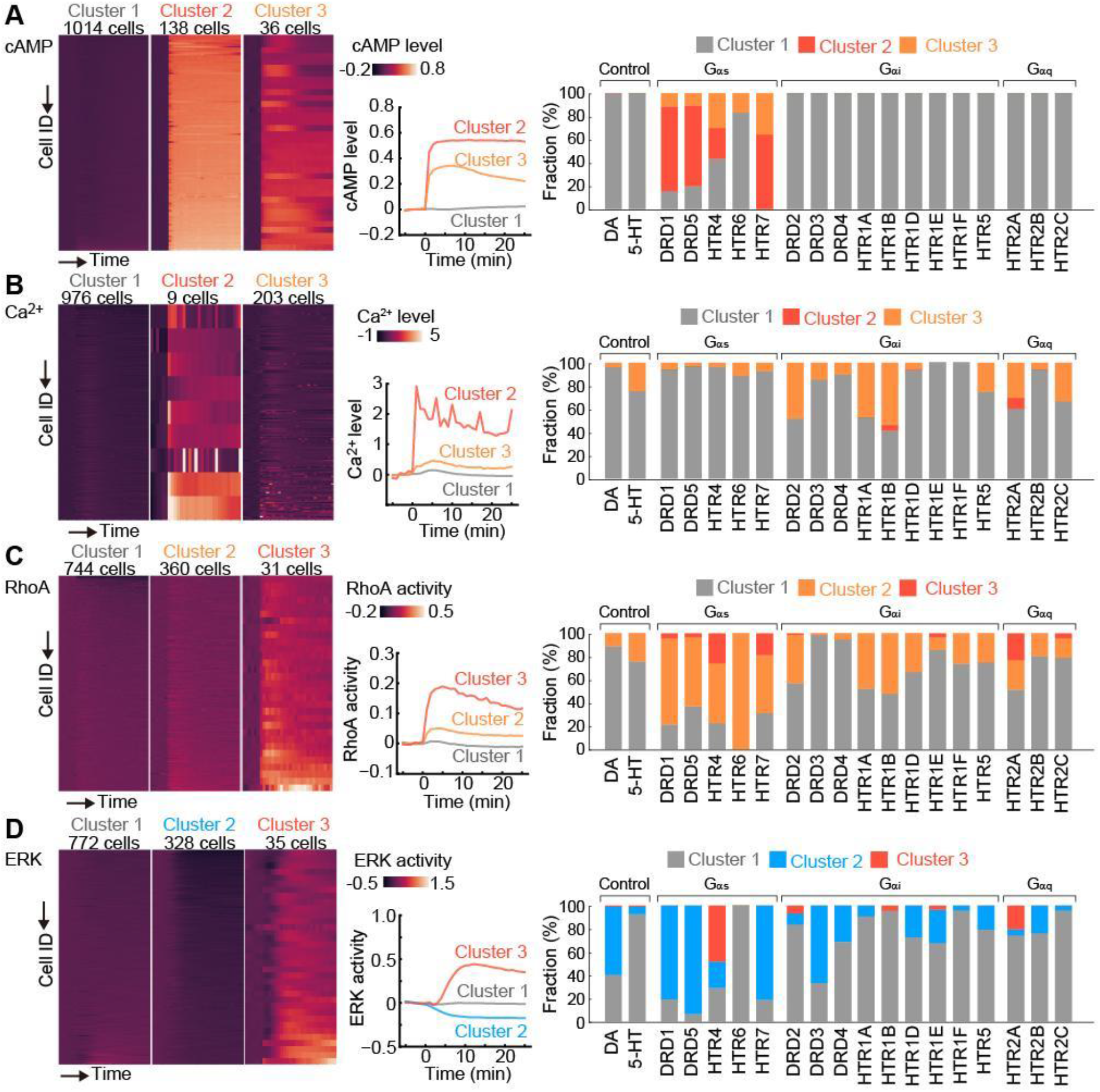
Systems analysis of signaling dynamics induced by activating dopamine or serotonin receptors. (A-D) Left heatmaps represent the result of time-series *K*-means clustering analysis. Vertical axis of heatmap represents each single cell and horizontal axis represents time (−5 min to 25 min since the ligand addition). Middle plots are the average time-course of each cluster. The proportion of clusters in each GPCR expressing cell are visualized in the right bar graphs. cAMP (A), Ca^2+^ (B), RhoA (C), and ERK (D), respectively.

Finally, we analyzed the interrelationships between GPCR downstream signaling dynamics. In our imaging system, two of four GPCR signaling are simultaneously observed in one cell, providing an opportunity to analyze crosstalk and feedback of two types of signal transduction at the single cell level. In addition to HeLa/cAMP/Ca^2+^ and HeLa/RhoA/ERK cell lines, we further established a stable cell line HeLa/cAMP/ERK expressing cAMP and ERK biosensors, thereby allowing to investigate the relations in cAMP-Ca^2+^, cAMP-ERK, and RhoA-ERK pairs. The dynamics of each signaling induced by activation of DRD1, HTR4, or HTR7 were visualized by heatmaps with sorting basal cAMP level or ERK activity (**Supplementary Figure S5**). Obviously, there is no correlation between cAMP and Ca^2+^ dynamics (**Supplementary Figure S5A**), consistent with the result that the activation of Gα_s_-coupled GPCR did not induce any changes in Ca^2+^ levels (**Figure 7A** and **7B**). ERK activation was often observed in HTR4 expressing cells with higher basal cAMP levels (**Supplementary Figure S5B**), whereas ERK activation seemed not to be correlated with RhoA activation (**Supplementary Figure S5C**). The findings provide a potential link between ERK and cAMP signaling, such as cross-talk regulations.

## DISCUSSION

In this study, we demonstrate the effectiveness of the live-cell fluorescence imaging system to simultaneously visualize multiple GPCR-induced downstream signal dynamics. Targeting dopamine receptors and serotonin receptors, we found that GPCR downstream signaling dynamics are highly variable. Although it is known which Gα protein the dopamine and serotonin receptors couple to, our experimental results indicate that the strength and temporal patterns of the downstream signaling vary among GPCRs. This is probably because the degree of Gα activation by GPCRs and/or inactivation of GPCRs via GRKs or β-arrestins differs among GPCRs. The crosstalk of downstream signaling of GPCRs may also affect their dynamics. In fact, the negative correlation between cAMP and ERK activity was observed at the single-cell level (**Supplementary Figure S5B**). This could be explained by the suppression of RAF by protein kinase A (PKA)[55,56]. In addition, the phosphodiesterases (PDEs) are regulated by Ca^2+^, PKA, and ERK [57,58], suggesting the potential contribution to the dynamics of these signaling molecules.

The single-cell imaging data further clarify the striking cellular heterogeneity of GPCR downstream signaling, showing diverse signaling dynamics such as responding vs. non-responding cells, sustained vs. transient responses, and pulsatile responses. The observed diversity of dynamics suggests that the GPCR signaling is enhanced by “dynamical encoding,” i.e., utilization of time courses of signaling activities as biological information. Despite dozens to hundreds of GPCRs being expressed in individual cells [59], GPCRs transmit signals primarily through only four types of Gα proteins, namely, Gα_s_, Gα_i/o_, Gα_q/11_, and Gα_12/13_. Thus, it is inevitable that multiple GPCRs in a single cell activate the same type of Gα proteins, making it difficult for cells to interpret the input information. The dynamical-encoding property has been found in many signaling components, including Ca^2+^ and ERK [60–62]. Pioneering works have revealed rich signaling dynamics downstream of GPCRs [63]. However, due to the limited number of GPCRs characterized in terms of dynamics, we are still far from unveiling the full diversity of GPCR signaling dynamics. For example, the cross-talk between the dynamics of cAMP and Ca^2+^ is of critical importance in neuromodulator signaling [64]. We thus considered that it would be of interest to apply our live-cell fluorescence imaging system to other neuromodulator GPCRs.

One unexpected finding was that heterodimerization of dopamine receptors did not enhance the intracellular Ca^2+^ increase induced by dopamine stimulation (**Supplementary Figure S2**). The most probable reason for this disparity in results is that we used different cell lines from the previous study (HeLa vs. HEK or neurons). A second possible reason is that the GPCR constructs we used in this study contained an HA signal peptide and FLAG-tag fused to the N-terminus and a V2-tail, TEV cleavage site, and rtTA fused to the C-terminus for the TANGO assay [18]. These moieties connected to the GPCRs might prevent the dopamine receptors from forming heterodimers. The third possibility is that the stoichiometry of the dopamine receptors was not appropriate. More careful analysis for GPCR heterodimers will be required in the future. Another unexpected finding was the constitutive activity of Gα_s_-coupled serotonin receptors, HTR4 and HTR6 (**Figure 6**). Constitutive activity has been reported in many types of GPCRs [65]. Indeed, it has been shown that HTR4 and HTR6 exhibit constitutive Gα_s_ activity [66–68]. The cellular heterogeneity of basal cAMP levels in cells expressing HTR4 or HTR6 is simply explained by the variability in the expression levels of those receptors and the constitutive activity of the receptors for Gα_s_. It would be interesting to investigate whether the expression levels of endogenous HTR4 and HTR6 are heterogenous, and if so, whether the cellular heterogeneity plays a physiological role.

We established a live-cell fluorescence imaging system for quantifying four GPCR downstream signals at the single-cell level. We would like to make two additional points in regard to technical matters. The first concern is endogenous GPCRs. In this study, we targeted dopamine and serotonin receptors, most of which are not expressed in HeLa cells, and therefore the data analysis was straightforward. However, in cases in which the endogenous GPCR responds to the ligand, the data should be analyzed with great care. The second concerns the biosensors for multiplexed fluorescence imaging. Because of the spectral overlap of biosensors, we established two types of reporter cell lines and co-cultured them, limiting multiplexed imaging at the single-cell level because of just two of the permutations for biosensor pairs. Ideally, it is desirable to establish one type of cell line expressing all reporters from the viewpoint of high-throughput screening. This issue could be solved by using the recently developed single fluorophore-based biosensors [71,72], making it easier to understand cross-talk and feedback control between signaling molecules at the single-cell level. In addition, recent advances in fluorescence imaging allow us to visualize multiple cell signaling at the single-cell level [73,74]. Further studies will be needed to visualize the downstream signaling dynamics of many types of GPCRs and reveal the information processing mechanism.

## Data availability

Data and reagents are available upon request to the corresponding authors.

## Author contributions

R.T., Y.G., and K.A. designed the research. R.T., Y.K., and Y.G. performed all experiments and data analysis. R.T., Y.G., Y.K., and K.A. wrote the manuscript.

## Acknowledgments

We thank all members of the Aoki Laboratory for their helpful discussions and assistance.

