## Supplementary data for "Quantitative live-cell imaging of GPCR downstream signaling dynamics"

### Title

### Affiliations

**Supplementary Figure S1**

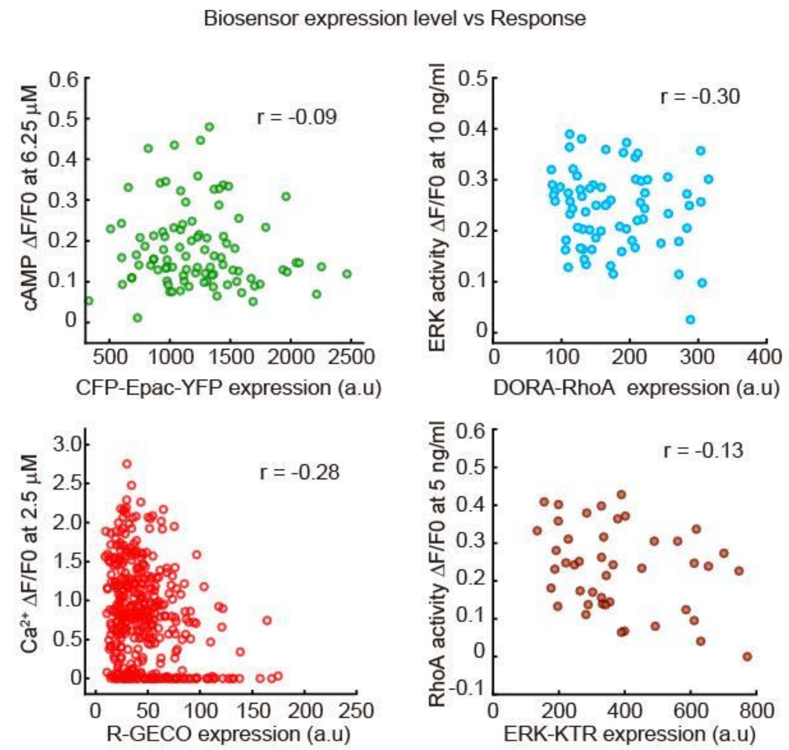


**Supplementary Figure S1. The expression level of biosensors does not correlate with their ligand-induced response.**

HeLa/cAMP/$Ca^{2+}$ cells and HeLa/RhoA/ERK cells were treated with submaximal concentration (nearby $EC_{50}$) of the control reagent used in Figure 2. The horizontal axis represents the biosensors' expression levels, whereas the vertical axis represents the response of each biosensor. Each dot represents the data from a single cell. The correlation coefficients (*r*) between the expression levels and responses are indicated in the right upper of graphs.

**Supplementary Figure S2**

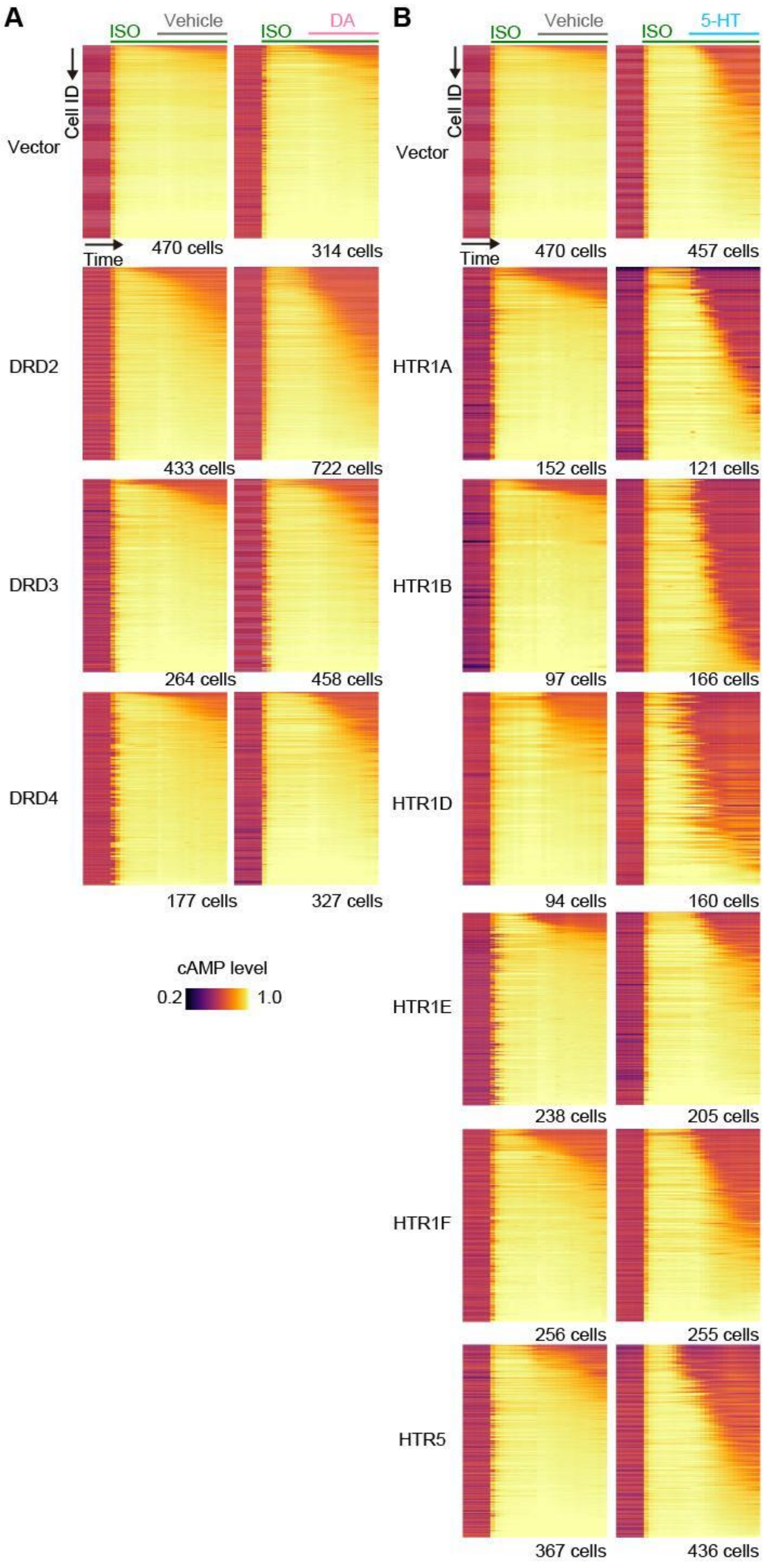

**Supplementary Figure S2. The dynamics of cAMP downregulation by G$\alpha_{i/o}$.**

(A and B) HeLa/cAMP/$Ca^{2+}$ cells expressing G$\alpha_{i/o}$-coupled dopamine receptors (A) or G$\alpha_{i/o}$-coupling serotonin receptors (B) were treated with 100 nM Isoproterenol (ISO) to increase the intracellular cAMP level through the activation of endogenous β-adrenergic receptors. Ten minutes after the ISO addition, cells were treated with vehicle, DA (A), or 5-HT (B). Normalized time-series of CFP/FRET of YFP-EPAC-CFP were visualized with a heatmap in which the vertical axis represents single cells, and the horizontal axis represents time (-5 min to 25 min since the ligand addition). Cells were sorted by the integrated CFP/FRET throughout the time course.

**Supplementary Figure S3**

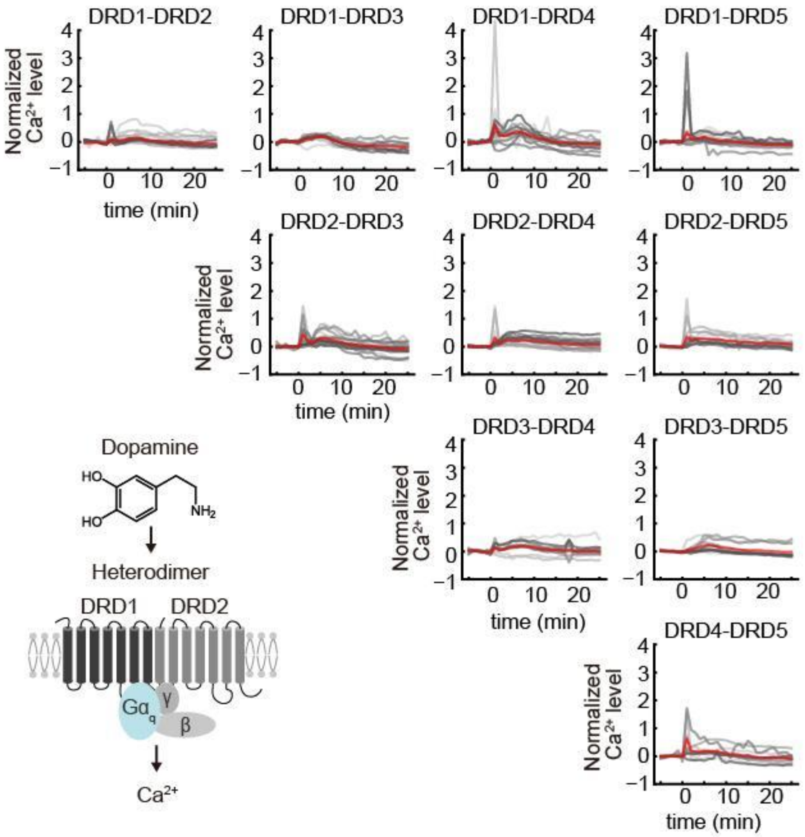


**Supplementary Figure S3. $Ca^{2+}$ dynamics of the coexpression of dopamine receptors upon dopamine stimulation.**

Normalized responses of $Ca^{2+}$ biosensors in HeLa cells expressing the indicated combination of dopamine receptors are shown. The fluorescent intensities of $Ca^{2+}$ biosensors are normalized by dividing by the averaged value before stimulation and plotted as a function of elapsed time after 10 μM dopamine administration. The red and gray lines represent the average and individual cells (n > 10 cells), respectively.

**Supplementary Figure S4**

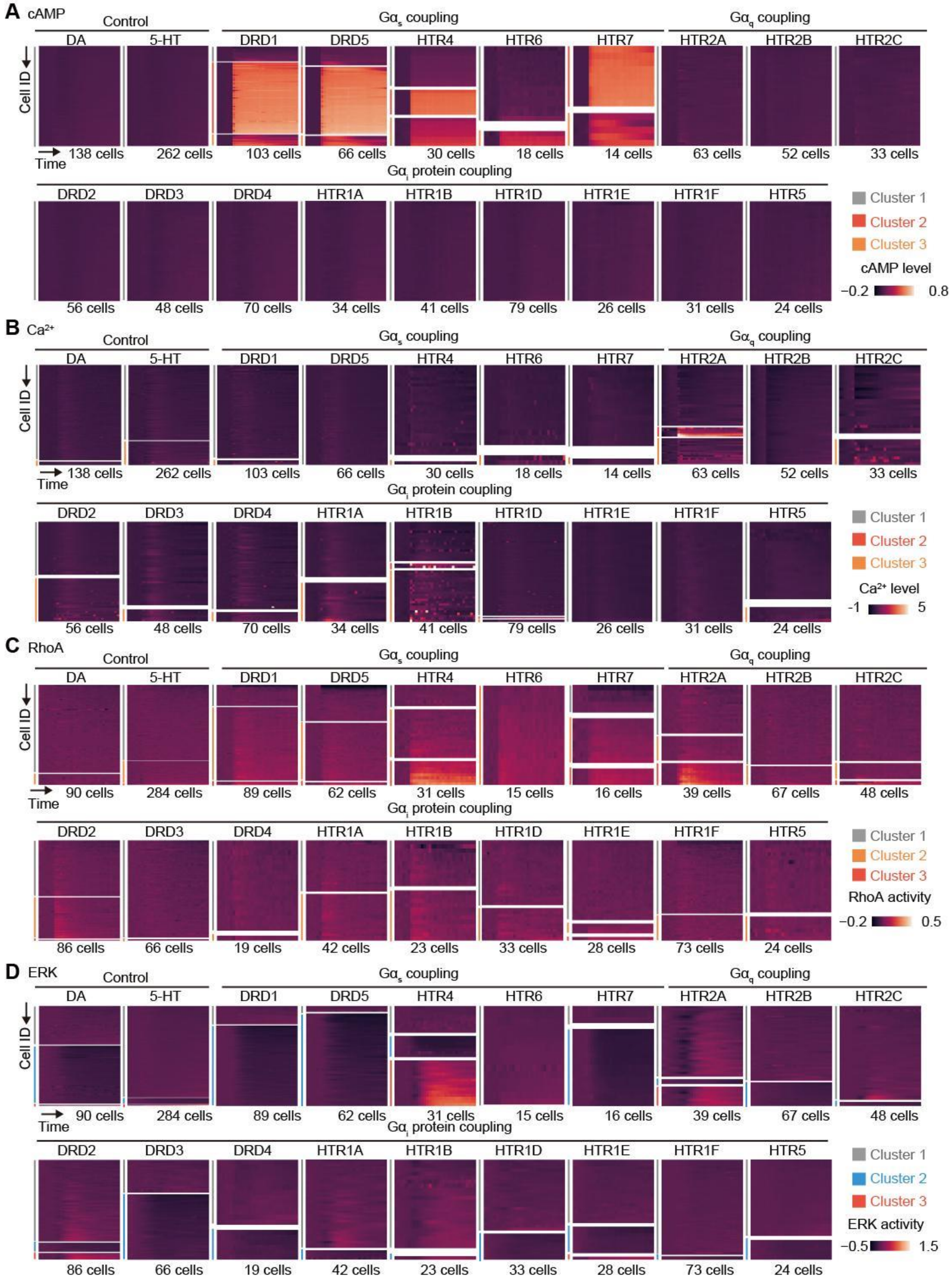

**Supplementary Figure S4. Classified all time-course single cell data of each receptors.**

Normalized single cell time-course data of cAMP (A), $Ca^{2+}$ (B), RhoA (C), and ERK (D) are subjected to the clustering analysis (see Materials and Methods) and represented as heatmaps. The vertical axis of heatmap represents single cells, and the horizontal axis represents time (-5 min to 25 min since the ligand addition).

**Supplementary Figure S5**

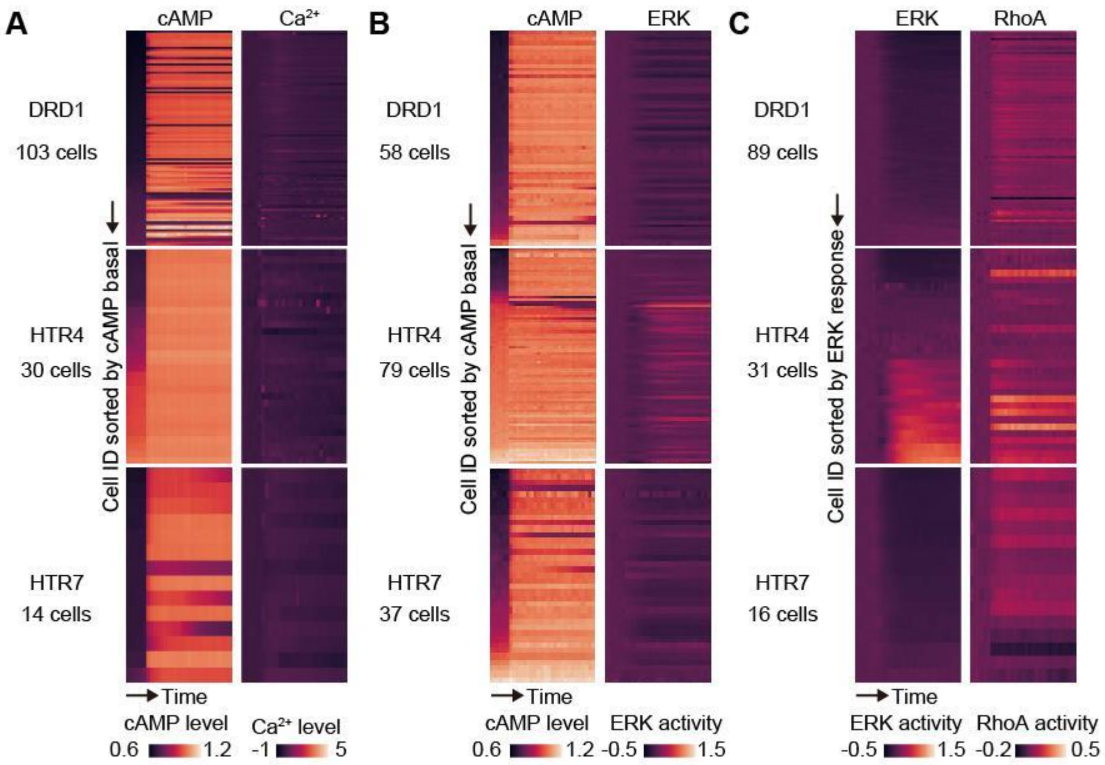


**Supplementary Figure S5. Multiplexed analysis of GPCR signaling.**

Time-course data from the same single-cell are shown side-by-side with heatmaps, in which the vertical axis represents single cells, and the horizontal axis represents time (-5 min to 25 min since the ligand addition). (A) Correlation of cAMP and $Ca^{2+}$. Cells were sorted by the basal cAMP levels. Note that only $Ca^{2+}$ response was normalized because of the heterogenous basal cAMP among HTR4 expressing cells. (B) Correlation of cAMP and ERK. Cells were sorted by the basal cAMP levels. $Ca^{2+}$ response was normalized. (C) Correlation of ERK and RhoA. Cells were sorted by the normalized ERK response.
